# Ancestry-Specific Predisposing Germline Variants in Cancer

**DOI:** 10.1101/2020.04.14.032557

**Authors:** Ninad Oak, Andrew D. Cherniack, R. Jay Mashl, TCGA Analysis Network, Fred R. Hirsch, Li Ding, Rameen Beroukhim, Zeynep H. Gümüş, Sharon E. Plon, Kuan-lin Huang

## Abstract

**Background:** Cancer risk differs across ancestries and these differences may result from differing prevalence of inherited genetic predisposition. Yet, most germline genomic studies performed to date have focused on individuals of European ancestry. Ancestry-specific analyses of germline genomes are required to inform cancer genetic risk and prognosis for each ancestral group. Here, we investigate potentially germline pathogenic variants in cancer predisposition genes (CPG) and their somatic effects in patients across diverse ancestral backgrounds.

**Methods:** We performed a retrospective analysis of germline genomic data of 9,899 patients from 33 cancer types generated by The Cancer Genome Atlas (TCGA) project along with matching somatic genomic and transcriptomic data. By collapsing pathogenic and likely pathogenic variants to the gene level, we analyzed the association between variants in CPGs and cancer types within each ancestry. We also identified ancestry-specific predisposing variants and their associated somatic two-hit events and gene expression levels.

**Results:** Recent genetic ancestry analysis classified the cohort of 9,899 cancer cases into individuals of primarily European, (N = 8,184, 82.7%), African (N = 966, 9.8%), East Asian (N = 649, 6.6%), South Asian (N=48, 0.5%), Native/Latin American (N=41, 0.4%), and admixed (N=11, 0.1%) ancestries. In the African ancestry, we discovered a potentially novel association of *BRCA2* in lung squamous cell carcinoma (OR = 41.4 [95% CI, 6.1-275.6]; FDR = 0.002) along with the previously identified association of *BRCA2* in ovarian serous cystadenocarcinoma (OR=8.5 [95% CI, 1.5-47.4]; FDR=0.045). Similarly, in the East Asian ancestry, we discovered one previously known association of *BRIP1* in stomach adenocarcinoma (OR=12.8 [95% CI, 1.8-90.84]; FDR=0.038). Rare variant burden analysis further identified 7 suggestive associations for cancer-gene pairs in African ancestry individuals previously well described in European ancestry including *SDHB* in pheochromocytoma and paraganglioma, *ATM* in prostate adenocarcinoma, *VHL* in kidney renal clear cell carcinoma, *FH* in kidney renal papillary cell carcinoma, and *PTEN* in uterine corpus endometrial carcinoma. Loss of heterozygosity was identified for 7 out of the 15 African ancestry carriers of predisposing variants. Further, tumors from the *SDHB* or *BRCA2* carriers showed simultaneous allelic specific expression and low gene expression of their respective affected genes; and *FH* splice-site variant carriers showed mis-splicing of *FH*.

**Conclusion:** While several predisposing genes are shared across patients, many pathogenic variants are found to be ancestry-specific and trigger somatic effects. Analysis of larger diverse ancestries genomic cohorts are required to pinpoint ancestry-specific genetic predisposition to inform personalized diagnosis and screening strategies.

## Background

Cancer risk differs across ancestries. According to the National Cancer Institute’s Surveillance, Epidemiology, and End Results (SEER) program, the cancer incidence per 100,000 ranges from 449 in race/ethnicity population self-identified as Whites, 453 in Blacks, 298 in Asian/Pacific Islanders, 315 in American Indian/Alaskan Natives, and 336 in Hispanics in the US between 2011-2015 [1]. While some of these differences may be attributed to non-genetic factors such as access to health care or diet, much can likely be explained by differences in the genomic architecture of these ancestries and differing frequencies of inherited genetic predisposition. Previous studies revealed different carrier rates of pathogenic variants across ancestries, albeit often in a limited panel of genes or selected cancer types[2–4].

While multiple large-scale genome-wide association studies have investigated the common risk variants contributing to cancer, fewer studies have interrogated rare pathogenic variants in non-European ancestries [5–9]. A 2019 systematic review of cancer sequencing studies found a total of only 764 reported non-European (minority) cases in 27 published studies with reported race/ethnicity [8]. Consequently, germline genetic testing in non-white patients often results in higher rates of variants of unknown significance (VUSs)[10]. Ongoing efforts are bridging the knowledge gap of cancer genetic predisposition in under-studied populations. Meanwhile, systematic cross-ancestry investigations of predisposing variants across cancer types are urgently needed to inform genetic testing for each ancestral group.

Herein, we analyzed germline variant data of 9,899 cancer cases across 33 cancer types from the Cancer Genome Atlas Project (TCGA) [11] to identify ancestry-specific cancer-gene associations where the gene is identified due to an excess of pathogenic/likely pathogenic germline variants the TCGA samples. In samples of African ancestry, we identified potentially two novel associations, *BRCA2* in lung squamous cell carcinoma (LUSC) and ovarian serous cystadenocarcinoma (OV). In analyses of individuals with East Asian ancestry, we identified one significant association for *BRIP1* in stomach adenocarcinoma (STAD). Using a complementary rare-variant association analysis, we identified seven additional suggestive cancer gene associations. Evidence of a somatic second hit event (i.e., loss of heterozygosity [LOH] or a biallelic mutation) was found in two-thirds of the tumors with germline predisposing variants. Many carriers of ancestry-specific predisposition variants showed altered expression of the affected genes, including elevated *RET* expression and reduced tumor suppressor gene expression compared to non-carriers, further supporting these genetic factors’ contribution to cancer predisposition.

## Methods

### Study Cohort and Genetic Ancestry Assignment

We used the clinical data provided by TCGA PanCanAtlas and restricted analyses to those with pass-QC blood/normal sequencing data. In addition to excluding cases with PanCanAtlas blacklisted germline BAM-files, cases with less than 60% genotype concordance between sequencing variant calls and SNP-genotype data were eliminated, where 10,389 cases were left. We further overlapped with the cases included in the PanCanAtlas AIM genetic ancestry assignment, resulting in the final set of 9,899 samples [11].

Genetic ancestry assignments were provided by the PanCanAtlas Ancestry Informative Markers (AIM) working group. The detailed descriptions of ancestry assignment procedures are available in the marker publication (**accepted draft attached**).

Briefly, consensus genetic ancestry for each TCGA case was determined as the majority of ancestry assignments that were independently determined by five methods across four institutions. These methods include those based on SNP-array genotypes used by Broad Institute, University of California San Francisco (UCSF), and Washington University (WashU), as well as those based on whole-exome sequencing data used by University of Trento and ExAC/Broad Institute. The five methods conducted variations of principal component analyses (PCA) on TCGA normal samples to infer genetic ancestry. We further provide the PCA plots showing the alignment of the major PCs in the UCSF and WashU analyses with the AIM-group consensus genetic ancestry in **Supp. Figure 1**.

For each sample, the percentage of global ancestry of African, European, East Asian, Native/Latin American, and South Asian (k=5) was further estimated using ADMIXTURE [17] version 1.23 based on the common SNP markers (1000 genome MAF > 1%) in the Broad Institute analysis. Samples with the proportion of the secondary ancestry greater than 20% were considered as admixed samples (**Supp. Table 1**). Sensitivity analyses revealed increased power by including admix samples in this cohort. Thus, cases with admixed ancestry assignments were grouped to their nearest neighbors (e.g., afr_admix to afr) for downstream analyses.

### Pathogenic and likely pathogenic germline variant calls

We downloaded the overall and predisposing germline variant calls previously reported by the PanCanAtlas Germline Analyses Working Group (https://gdc.cancer.gov/about-data/publications/PanCanAtlas-Germline-AWG)[11]. The detailed description of variant calling and classification procedures are available in the TCGA PanCanAtlas germline publication [11].

Briefly, germline SNVs were identified using the union of variant calls between Varscan[12] and GATK[13]. Germline indels were identified using Varscan, GATK, and Pindel[14], and we only retained variants called by at least two out of the three callers or high-confidence Pindel-unique calls (at least 30x coverage and 20% VAF). We used the GRCh37-lite reference. We further required the variants to have an Allelic Depth (AD) ≥5 for the alternative allele. We then used bam-readcount to quantify the number of reference and alternative alleles in both normal and tumor samples. We required the variants to have at least 5 counts of the alternative allele and an alternative allele frequency of at least 20%. Of these, we included those rare variants with ≤0.05% allele frequency in 1000 Genomes and ExAC (release r0.3.1). We subsequently retained only cancer-relevant pathogenic variants, based on whether they were found in the curated cancer variant databases or a 152 curated cancer predisposing gene list. Finally, we manually reviewed all variants using IGV and filtered out variants with poor support sequence reads.

The variants defined by the above pipeline were then classified using an automatic pipeline termed CharGer [15] (https://github.com/ding-lab/CharGer) that adopts the American College of Medical Genetics and Genomics/Association of Molecular Pathology (ACMG/AMP) variant classification guidelines which are designed for assessment of germline variants in Mendelian disorders [16]. For the CharGer classification pipeline, we defined 12 pathogenic evidence levels and 4 benign evidence levels using a number of datasets, including ExAC and ClinVar. The pathogenic evidence adds points, whereas benign evidence subtracts points that amount to pathogenicity (pathogenic requires the variant to be described as pathogenic by the reviewed clinical significance in ClinVar (not including variants showing “Conflicting interpretations of pathogenicity”) or other cancer predisposition gene databases, likely pathogenic requires CharGer score > 8). To acquire enough CharGer points to be classified as likely pathogenic, the variants typically need to be predicted to result in truncation in cancer predisposition genes where loss of function (LOF) is a known disease mechanism and harbor variants with a dominant (evidence level PVS1, +8 points) or a recessive (evidence level PSC1, +4 points) mode of inheritance. Additionally, evidence level PS1, +7 points, are scored if the variant results in the same peptide sequence change as an established pathogenic variant. All other modules will each add ≤2 points.

### Principal Component Analysis (PCA)

Birdseed genotype files were downloaded from Genomic Data Commons (GDC) in the legacy (hg19) archive onto Institute for System Biology-Cancer Genome Cloud (ISB-CGC), converted to individual VCF files, and then merged into a combined VCFs containing 11,459 samples and 522,606 variants. We conducted PCA as implemented by PLINK (v1.9)[18]. Specifically, we retained 298,004 variants with MAF >0.15 for population structure analysis. The resulting eigen values and eigen vectors were then recorded. PC1 and PC2 accounted for 51.6% and 29.2% of the variations across the first 20 PCs, and none of the trailing PCs accounted for more than 3.2%. Thus, we subsequently controlled for PC1 and PC2 in the ancestry-specific cancer predisposing gene analysis (**Supp. Figure 1)**.

### Multivariate regression to identify the enrichment of pathogenic variants

For each cancer type within each ancestry, we conducted multivariate logistic regression analyses considering the case status of the cancer type as the dependent variable (using all other cancer cohorts as controls) and the carrier status of each predisposing gene as an independent variable. The model corrected for age at the initial pathologic diagnosis, gender, the first two principal components (accounted for 80.8% variations across the first 20 PCs). All ancestry cohorts are called using the same variant calling pipeline, thus avoiding the potential danger of comparing this population against other cohorts such as ExAC. We collapsed predisposing (pathogenic and likely pathogenic) germline variants to the gene level. Only ancestry-cancer combinations with at least 20 cases and predisposing genes with at least two individuals with predisposing variants within the cohort are tested. In total, we tested 33 cancers in European Ancestry, 15 cancers in African Ancestry, and 8 cancers in East Asian ancestry that met this criterion. No cohorts of the Native/Latin American and South Asian ancestry have sufficient sample sizes in TCGA for testing. Among these tested cancers, we tested a total of 114 cancer-gene combinations for multivariate regression analysis, of which 101 were within European ancestry, 9 were in African ancestry, and 4 were in East Asian ancestry. P values were calculated using the Wald test and adjusted to FDR using the standard Benjamini-Hochberg procedure.

### Burden testing of pathogenic variants

We conducted burden testing of the cohort within each ancestry as defined by the TCGA AIM working group. Specifically, we adopted the Total Frequency Test (TFT)[19] by collapsing predisposing (pathogenic and likely pathogenic) germline variants to the gene level. For each cancer type with at least 20 cases of the tested ancestry with at least one predisposing variant carrier, we tested the burden of predisposing variants for each gene against all other cancer cohorts as controls. Among the cancers that met the sample size criteria described above, we tested a total of 120 cancer-gene combinations using rare variant burden testing, of which 104 were within European ancestry, 11 were in African ancestry, and 5 were in East Asian ancestry. The resulting P values were adjusted to FDR using the standard Benjamini-Hochberg procedure.

### gnomAD analysis

We analyzed the gene-level and variant-level frequency of the identified genetic predisposition using the non-cancer subset of the genome aggregation database (gnomAD-non-cancer) cohort (118,479 WES and 15,708 WGS samples) [20,21] (http://gnomad.broadinstitute.org) [Date Accessed: February 2020]. For the gene-level analysis, we retained rare variants with ancestry-specific minor allele frequency <0. 5%. We further retained pathogenic and likely pathogenic variants per ACMG/AMP criteria as ascertained by InterVar [22] and annotated using ANNOVAR [23]. Allele frequencies were summarized at gene-level within each sub-population in gnomAD using total allele counts and maximum allele numbers within each group.

The lolliplot diagrams in Figure 2 were constructed and modified from the PCGP protein paint (https://pecan.stjude.cloud/proteinpaint) based on the specified RefSeq transcript.

### Expression Analysis

TCGA level-3 normalized RNA expression data were downloaded from Firehose (2016/1/28 analysis archive). The tumor expression percentile of individual genes in each cancer cohort was calculated using the empirical cumulative distribution function (ecdf), as implemented in R. We annotated germline carriers of predisposition variants with extreme mRNA tumor expression (>80^th^ or < 20^th^ percentile) of the affected gene. For samples within the same ancestry and same cancer cohort, we then used the two-sample Kolmogorov-Smirnov test to compare the expression percentile distribution between variants of oncogenes and tumor suppressors. The resulting P values were adjusted to FDR using the standard Benjamini-Hochberg procedure.

For the ancestry-specific variants, we recorded the RNA variant allele fraction (RNA VAF) of the mutant allele in the RNA-Seq bam files. For splice site variants, we assessed the mis-splicing of the transcript and variants using IGV.

### Power and Down-sampling Analysis

*Post hoc* power analyses were performed using R-package SKAT[24] and the power_logistic function to calculate the number of samples for rare variant association with causal percentage = 80%, minor allele frequency < 0.1%, and using OR >1 through OR<10. Each calculation was performed using 100 simulations over a target 5kb region.

Additionally, we performed a down-sampling analysis for each tumor type by random sampling of subsets of samples with incremental sizes from zero to the total number of samples in that tumor type. We identified the number of significantly mutated genes as described above within each subset and plotted a smoothed function (loess method) against the subset size. Each calculation was performed at ten iterations **(Supp. Figure 2)**.

## RESULTS

### Ancestry Demographics of TCGA Cohort

We classified the 9,899 TCGA cases with pass-QC germline data across 33 cancer types by genotype-defined ancestries defined by the PanCanAtlas Ancestry Informative Markers (AIM) working group (**Supp. Figure 1**, **Methods**, Table 1**Error! Reference source not found.**). The European ancestry contained 82.68% (N=8,184) of individuals in this cohort. The remainder of the cohort consisted of 9.76% (n=966) African ancestry, 6.56% (n=649) East Asian ancestry, 0.48% (n=48) South Asian ancestry, 0.41% (n=41) Native/Latin American ancestry, and 0.11% (n=11) mixed ancestry. The largest ancestry-specific tumor cohorts are breast invasive carcinoma (BRCA) for the European ancestry (n=811) and African ancestry (n=180), liver hepatocellular carcinoma (LIHC) for the East Asian ancestry (n=162), and thyroid carcinoma (THCA) for the Native/Latin American ancestry (n=11) and the South Asian ancestry (n=11).

**Table 1.** The demographic distribution of TCGA PanCancerAtlas cohort

|  | Cancer | European | African | Native/Latin American | East Asian | South Asian | Admix | Total |
| --- | --- | --- | --- | --- | --- | --- | --- | --- |
| Total Number (%) |  | 8184 (82.68%) | 966 (9.76%) | 41 (0.41%) | 649 (6.56%) | 48 (0.48%) | 11 (0.11%) | 9899 (100%) |
| ACC | Adrenocortical Carcinoma | 82 | 2 | 0 | 2 | 0 | 0 | 86 |
| BLCA | Bladder Urothelial Carcinoma | 329 | 21 | 1 | 43 | 2 | 0 | 396 |
| BRCA | Breast Invasive Carcinoma | 811 | 180 | 5 | 53 | 8 | 1 | 1058 |
| CESC | Cervical Squamous Cell Carcinoma and Endocervical Adenocarcinoma | 208 | 31 | 1 | 22 | 0 | 0 | 262 |
| CHOL | Cholangiocarcinoma | 30 | 2 | 0 | 2 | 0 | 0 | 34 |
| COAD | Colon Adenocarcinoma | 343 | 58 | 0 | 12 | 0 | 0 | 413 |
| DLBC | Lymphoid Neoplasm Diffuse Large B-cell Lymphoma | 23 | 0 | 0 | 15 | 1 | 0 | 39 |
| ESCA | Esophageal Carcinoma | 126 | 4 | 0 | 44 | 0 | 0 | 174 |
| GBM | Glioblastoma Multiforme | 292 | 37 | 0 | 2 | 1 | 0 | 332 |
| HNSC | Head and Neck Squamous Cell Carcinoma | 439 | 50 | 6 | 7 | 5 | 1 | 508 |
| KICH | Kidney Chromophobe | 56 | 4 | 0 | 1 | 1 | 0 | 62 |
| KIRC | Kidney Renal Clear Cell Carcinoma | 306 | 54 | 2 | 7 | 1 | 0 | 370 |
| KIRP | Kidney Renal Papillary Cell Carcinoma | 207 | 63 | 0 | 6 | 1 | 0 | 277 |
| LAML | Acute Myeloid Leukemia | 125 | 14 | 0 | 2 | 0 | 0 | 141 |
| LGG | Brain Lower Grade Glioma | 455 | 23 | 4 | 10 | 3 | 0 | 495 |
| LIHC | Liver Hepatocellular Carcinoma | 179 | 18 | 0 | 162 | 1 | 1 | 361 |
| LUAD | Lung Adenocarcinoma | 416 | 52 | 1 | 9 | 0 | 0 | 478 |
| LUSC | Lung Squamous Cell Carcinoma | 455 | 29 | 0 | 11 | 0 | 0 | 495 |
| MESO | Mesothelioma | 78 | 0 | 0 | 0 | 1 | 0 | 79 |
| OV | Ovarian Serous Cystadenocarcinoma | 348 | 30 | 1 | 7 | 5 | 0 | 391 |
| PAAD | Pancreatic Adenocarcinoma | 158 | 9 | 0 | 11 | 0 | 1 | 179 |
| PCPG | Pheochromocytoma and Paraganglioma | 146 | 20 | 0 | 3 | 4 | 0 | 173 |
| PRAD | Prostate Adenocarcinoma | 411 | 59 | 0 | 9 | 2 | 1 | 482 |
| READ | Rectum Adenocarcinoma | 135 | 6 | 0 | 1 | 0 | 0 | 142 |
| SARC | Sarcoma | 217 | 18 | 0 | 6 | 0 | 2 | 243 |
| SKCM | Skin Cutaneous Melanoma | 448 | 1 | 2 | 12 | 0 | 0 | 463 |
| STAD | Stomach Adenocarcinoma | 294 | 15 | 0 | 90 | 0 | 0 | 399 |
| TGCT | Testicular Germ Cell Tumors | 107 | 4 | 0 | 3 | 0 | 0 | 114 |
| THCA | Thyroid Carcinoma | 359 | 33 | 11 | 52 | 11 | 3 | 469 |
| THYM | Thymoma | 96 | 8 | 0 | 12 | 0 | 0 | 116 |
| UCEC | Uterine Corpus Endometrial Carcinoma | 382 | 112 | 7 | 30 | 1 | 1 | 533 |
| UCS | Uterine Carcinosarcoma | 43 | 9 | 0 | 3 | 0 | 0 | 55 |
| UVM | Uveal Melanoma | 80 | 0 | 0 | 0 | 0 | 0 | 80 |

### Ancestry-Specific Cancer Predisposing Genes

Acknowledging the limited power to assess ancestry-specific associations as shown by the *Post Hoc* power analyses (**Supp. Figure 2**), we sought to identify cancer predisposing genes within each ancestry. We considered cancer predisposing genes as those statistically enriched for pooled pathogenic and likely pathogenic variants (referred to here as predisposing variants) as previously classified[11]). For each ancestry-cancer type pair, we conducted multivariate regression analyses correcting for onset age, gender, and the first two principal components.

Along with 36 cancer-gene associations (FDR <0.05, Wald test) found in the European ancestry, we identified two specific cancer-gene associations in the African ancestry: *BRCA2* in ovarian cancer (OV) (OR=8.5 [95% CI, 1.5-47.4]; FDR=0.045) and LUSC (OR = 41.4 [95% CI, 6.1-275.6]; FDR = 0.002). We also identified one association in the East Asian ancestry, *BRIP1* in STAD (OR=12.8 [95% CI, 1.8-90.84]; FDR=0.038) (**Figure 1, Supp. Table 2a**). While the association of BRCA2 and LUSC is first described in African-American ancestry here, *BRCA2* was also recently found to be associated with LUSC in the European ancestry [25]. The association of *BRIP1* predisposition to STAD in the East Asian ancestry was also previously reported for the European ancestry [26]. These findings (including novel associations) in a large heterogeneous cancer population build on older studies that evaluated individual cancer predisposition genes and cancer risk across ancestries.

The top associated predisposing genes and their carrier frequency vary widely across ancestries (Figure 1A). For genes with a significant association in the African ancestry, we observed a higher carrier frequency compared to other ancestries. For example, in LUSC, *BRCA2* predisposing variants were found in 2 of the 29 African ancestry samples (6.9%), whereas we only found 1 *BRCA2* carrier out of the 455 European-ancestry samples (0.44%).

**Figure 1.**
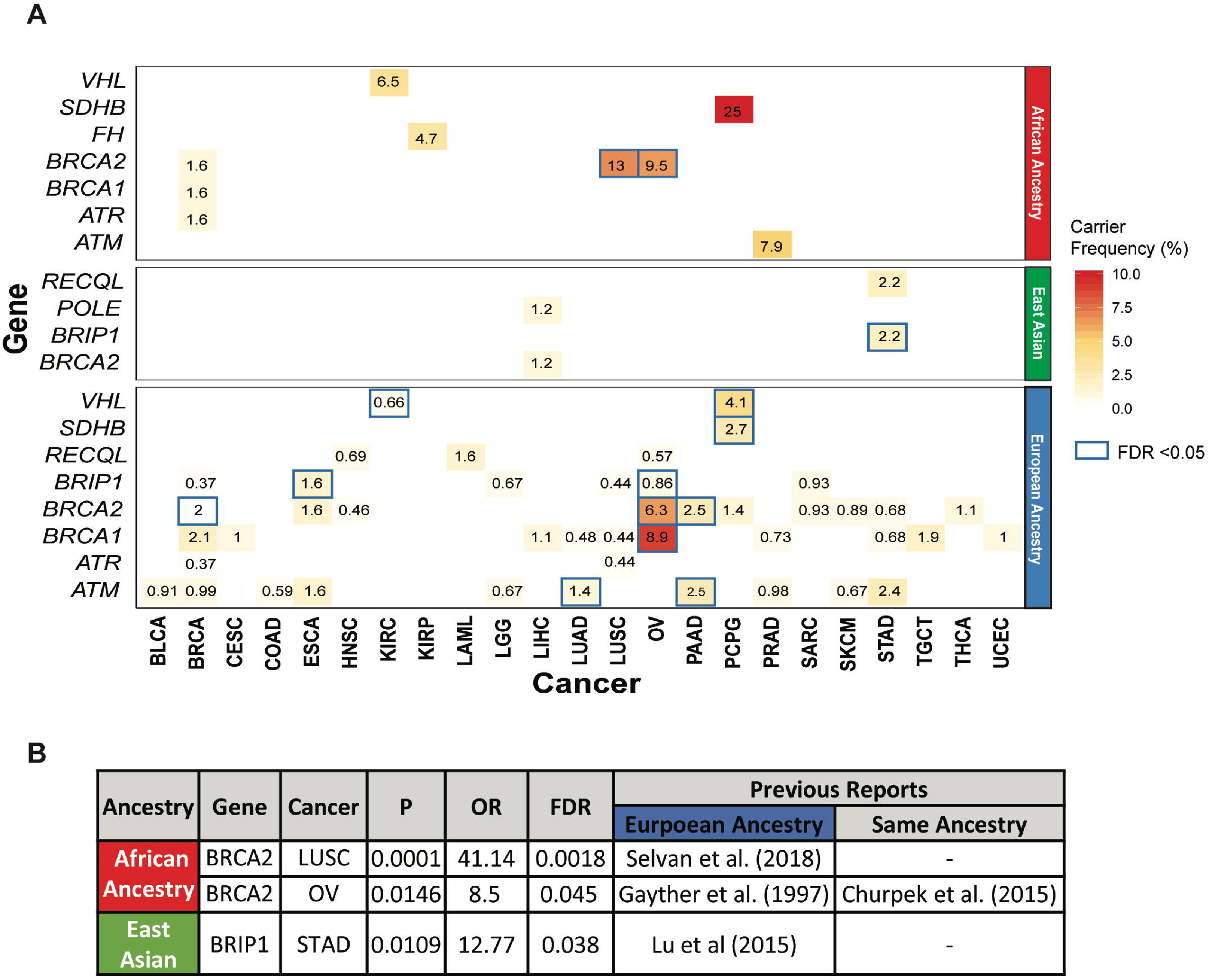
Cancer predisposing genes identified in each ancestry across 9,899 TCGA cases across cancer types in the African ancestry, East Asian, and European ancestries. **A.** Ancestry-specific cancer-gene pairs from TCGA dataset containing cancer predisposing variants as identified by multivariate logistic regression analyses. Each number represents carrier frequencies of predisposing genes within that cancer cohort. Genes with significant associations (Wald test FDR < 0.05) are highlighted with blue boxes. **B.** Significant cancer-predisposing gene associations (FDR < 0.05) identified in the African and East Asian ancestries.

We next investigated whether the cross-ancestry differences in predisposing gene frequencies were also observed in other cohorts. Specifically, we examined the gene-level rates of individuals carrying pathogenic and likely pathogenic variants in the gnomAD non-cancer cohort [20,21] (118,479 WES and 15,708 WGS samples, **Methods, Supp. Table 3**). *BRCA2* showed the highest frequency in the African ancestry (0.072%) than all other defined ancestries, including non-Finnish European (0.048%) and East Asian (0.047%). *BRIP1* also showed higher frequency in the East Asian ancestry (0.068%) than all ancestries (≤0.045%) except for the non-Finnish European ancestry (0.099%).

To generate hypotheses for future targeted studies, we investigated additional ancestry- implicated genes using total frequency testing (TFT) of predisposing variants, fully acknowledging potential confounders using this method (**Supp. Table 2b**). We identified 7 suggestive (FDR < 0.05 in the TFT analysis) ancestry-specific cancer-gene associations in the African ancestry, 6 of which have been previously described including *SDHB* in PCPG[27], *ATM* in PRAD[28,29], *FH* in KIRP[30], *VHL* in KIRC[31], *PTEN* in UCEC[32], and *BRCA2* in OV[33]. We also re-discovered the potentially novel association of *BRCA2* in LUSC described above. In the East Asian ancestry, we identified 3 borderline-suggestive associations (FDR = 0.32): *RECQL* in STAD, *BRIP1* in STAD, and *POLE* in LIHC. In STAD, *RECQL* and *BRIP1* each affected 2 of the 90 East Asian ancestry cases, but none of the 294 European-ancestry cases. In LIHC, two protein-truncating variants were seen in *POLE* among 162 East Asian ancestry cases compared to none in 179 European-ancestry cases. These suggestive associations remain to be established and are only used to identify potential predisposing variants with supporting somatic evidence.

### Ancestry-Specific Predisposing Variants

We next examined ancestry-specific predisposition at the variant level (Figure 2, **Supp. Table 4**) for the 3 significant associations from the multivariate logistic regression analyses and the 7 suggestive associations from the TFT analysis. The cancer-gene pairs included 15 predisposing variants within the African ancestry and another 6 within the East Asian ancestry.

Notably, none of the above variants discovered in the African ancestry were observed in any other ancestry within that cancer type (Figure 2). Across the pan-cancer TCGA cohort, all of the *BRCA2* frameshift variants found in LUSC and OV were unique to the African ancestry. For other associated genes in the African ancestry, including *ATM* (PRAD)*, FH* (KIRP), and *VHL* (KIRC), the predisposing variants differ between the African and European ancestries (Figure 2B). The African-ancestry-specific predisposing variants include splice site variants *ATM* c.2921+1G>A and *FH* c.556-2A>T, protein-truncating variants *ATM* p.T2333fs and *FH* p.S187*, and missense variants *ATM* p.R3008C. *VHL* p.C162F is the only recurrent variant found in two KIRC cases.

**Figure 2.**
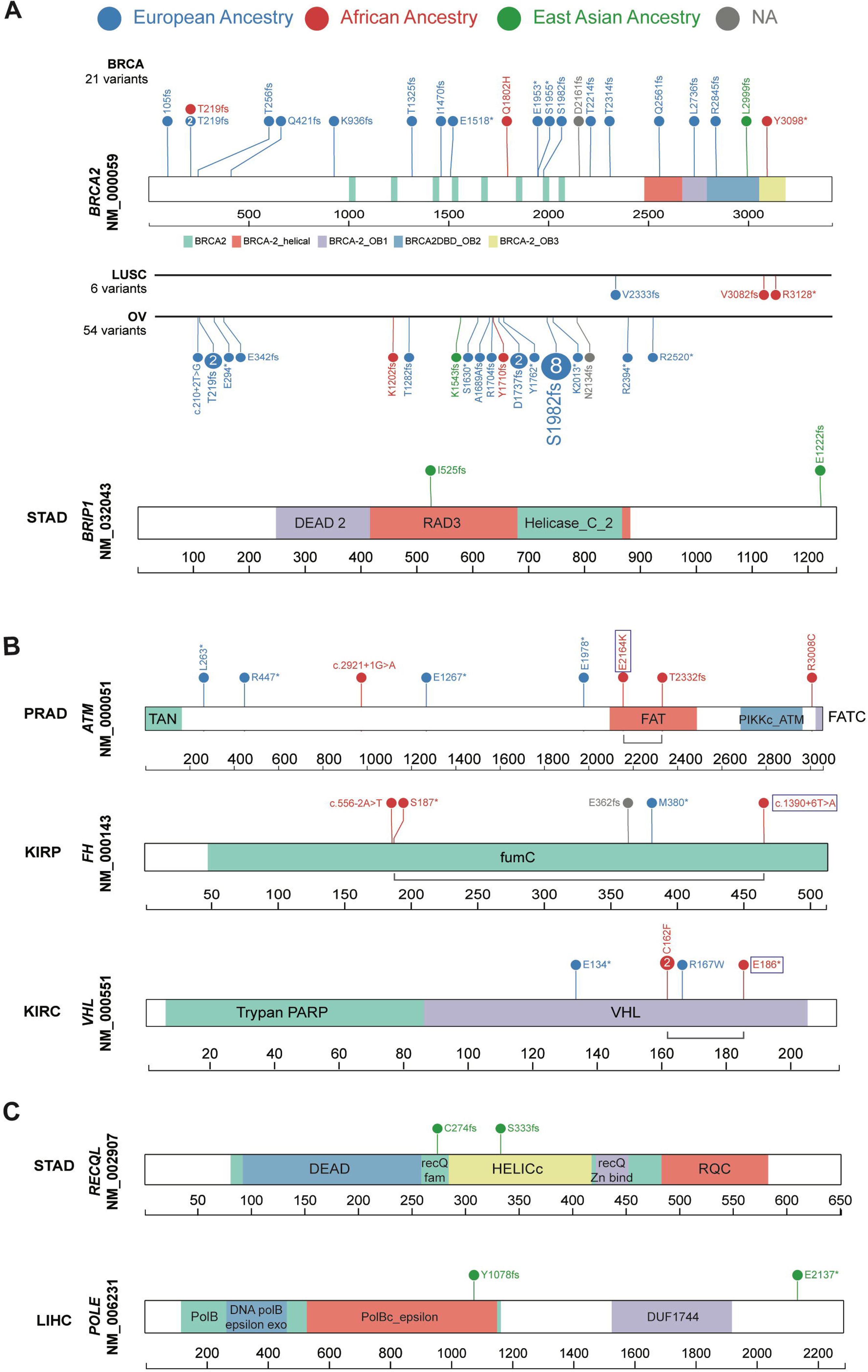
Ancestry-specific predisposing germline variants. Predisposing variants in the significant (regression analysis) and suggestive (rare variant burden testing) cancer-gene associations are shown. The variants are labeled with carrier counts and colored by their respective carriers’ ancestry (European: blue, African ancestry: red, East Asian: green). **A.** Predisposing variants identified in the African and East Asian ancestries are shown across respective cancer types. For *BRCA2*, predisposing variants across all cancers are shown (top) in comparison with the two cancer types with significant associations in the African ancestry (LUSC and OV, bottom). Similarly, predisposing variants contributing to the significant association of *BRIP1* in STAD in the East Asian ancestry, are shown. **B.** Suggestive predisposing variants identified in the African ancestry are shown for *ATM*, *FH*, and *VHL* genes within their associated cancer types. Bi-allelic events in each carrier are linked by a grey line bracket where the somatic second-hit mutations are marked with a box. **C.** Borderline-suggestive predisposing variants identified in the East Asian ancestry are shown for *RECQL* in STAD and *POLE* in LIHC.

In the East Asian ancestry, we assessed predisposing variants in *BRIP1* (STAD), *POLE* (LIHC), and *RECQL* (STAD) (Figure 2A and C). These include two *BRIP1* variants p.I525fs and p.E1222fs and two protein-truncating variants in *POLE* and *RECQL*, respectively. All six predisposing variants were not shared with any other ancestry in the TCGA cohort (Figure 2C).

To further investigate the carrier frequencies of the predisposing variants across ancestries, we analyzed the frequency of these variants of the gnomAD non-cancer dataset [20,21]. Among the African ancestry-specific predisposing variants, splice-site variant *ATM* c.2921+1G>A (African ancestry Allelic Count [AC]/ Total Allele Number [AN]=1/14,878; Allelic Frequency [AF] = 0.0067%) and *BRCA2* p.R3128* (African ancestry AC/AN = 4/23,610; AF = 0.016%) were the only variants present in the African and non-Finnish European ancestries in gnomAD-non-cancer dataset. All other variants were absent within African ancestry and most other ancestries in gnomAD except *SDHB* p.R46X (Finnish European ancestry AC/AN = 2/25,066; AF = 0.007%) and *ATM* p.R3008C (East Asian ancestry AC/AN = 1/17,688; AF = 0.005%). Similarly, only two of the six East Asian ancestry-specific predisposing variants, *BRIP1* p.E1222Gfs (East Asian ancestry AC/AN = 11/19,232; AF = 0.05%) and *POLE* p.Tyr1078fs (East Asian ancestry AC/AN = 1/17,692; AF = 0.005%), were present exclusively in the East Asian ancestry of gnomAD-non-cancer dataset. Of note, 7 of the 15 predisposing variants, including *BRCA2* variants in OV (p.Y1710fs, p.K1202fs) and in LUSC (p.V3082fs), were not found in ClinVar [34]. While *VHL* p.C162F lacks a ClinVar record, the co-localizing p.C162W showed three reports of pathogenicity and one report of uncertain significance.

We also investigated the presence of the six predisposing variants in the East Asian ancestry from the gnomAD non-cancer dataset. Only the *POLE* p.Y1078fs (AC/AN=1/17,692, AF= 0.0056%) and *BRIP1* p.E1222fs (AC/AN=11/19,232, AF=0.057%) were present exclusively in the East Asian ancestry of gnomAD-non-cancer dataset. All other East Asian-ancestry variants were not detected in this dataset. Of note, none of the six variants were previously reported in ClinVar[34].

### Germline-somatic Two-hit Events

We next examined the two-hit hypothesis, whereby a somatic second hit of the same gene is found in carriers of the germline predisposing variants [35,36]. First, we investigated the extent of loss of heterozygosity (LOH) of the predisposing variants using our previously developed statistical test [26] (**Methods**) that compares the variant allele fractions in tumor vs. normal samples. Among the variants observed in the African ancestry, we observed significant LOH (FDR <0.05) for both truncating variants in *SDHB* p.R116fs and p.R46* in PCPG (Figure 3A). Three additional variants exhibited significant LOH, including *BRCA2* p.R3128* (LUSC), *BRCA2* p.K1202fs (OV), and *FH* p.S187* (KIRP). We also observed suggestive LOH (FDR < 0.15 or tumor VAF > 0.6) for *ATM* c.2921+1G>A (PRAD) and *BRCA2* p.Y1710fs (OV) (Figure 3B). Among the six predisposing variants in the East Asian ancestry, only *POLE* p.E2137* (LIHC) showed significant LOH (Figure 3A).

**Figure 3.**
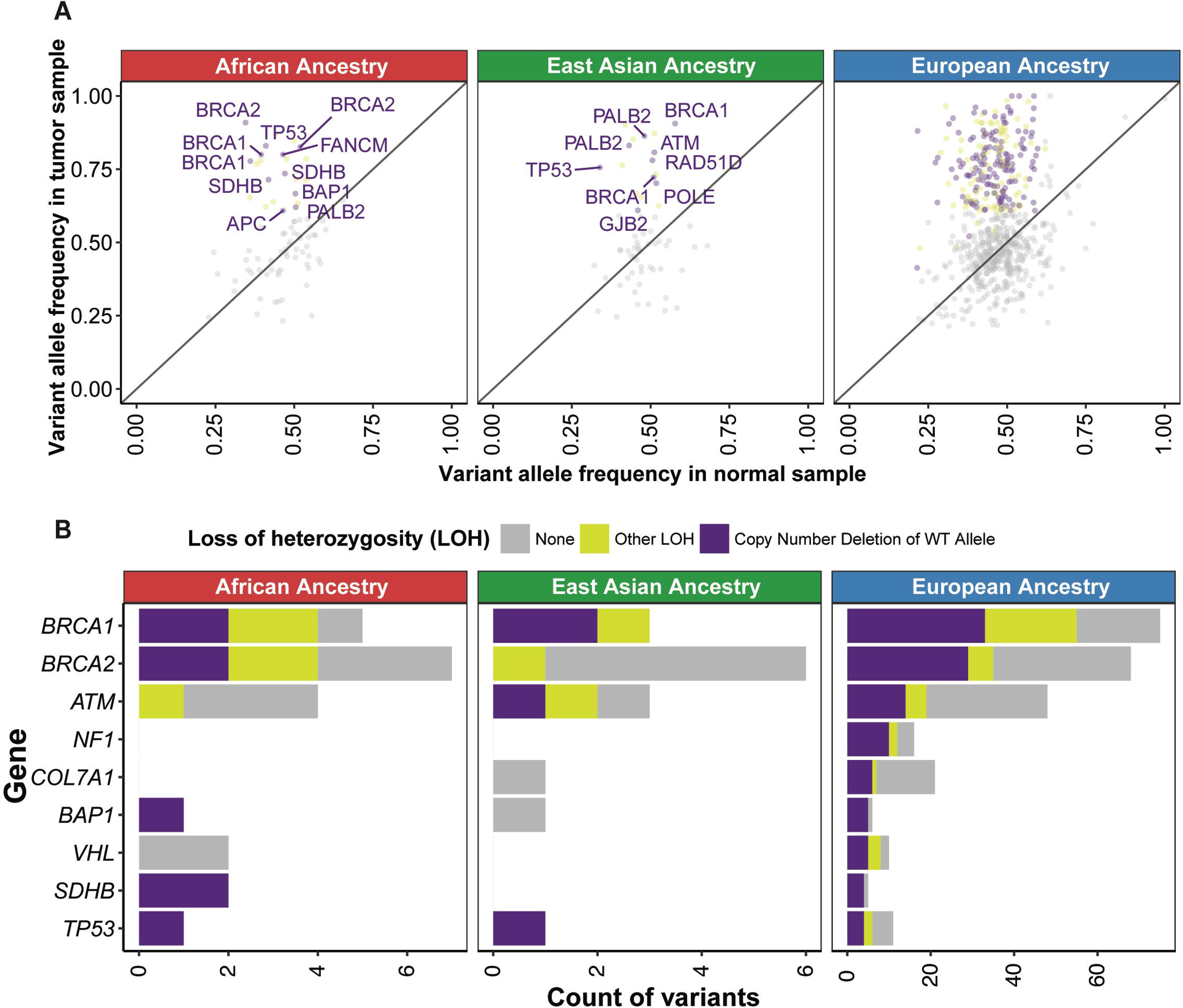
Loss of heterozygosity (LOH) and transcriptional effects associated with ancestry-specific predisposing germline variants. **A.** LOH in ancestry-specific predisposing variants shown by comparing variant allele frequency in tumor vs. that in normal samples. Each dot denotes a variant and the affected genes are labeled in cases where showed both significant allelic imbalance and copy number deletion of the wild-type alleles (in purple). Variants showing significant allelic imbalance yet no conclusive evidence of wild-type alleles are considered as other LOH and marked in yellow. All other variants are shown in grey. **B.** Count distribution of each type of LOH events across genes in the African ancestry, the East Asian ancestry, and the European ancestry. Note given the larger number of events, the x-axis for the European ancestry is shown on a different scale.

As an alternative mechanism of a somatic second-hit, we identified three biallelic mutations where the rare germline predisposing variant was coupled with a second somatic mutation of the same gene, all found in African ancestry carriers (labeled in Figure 2B, **Supp. Table 4b**). In a PRAD carrier of *ATM*, the germline p.L2332fs variant was coupled with a somatic p.E2164K mutation; in the KIRC carrier of *VHL*, the germline p.C162F variant was coupled with somatic p.E186* mutation and in a KIRP carrier of *FH*, germline p.S187* variant was coupled with a somatic splice-site mutation c.1390+6T>A. Analysis of RNA from the KIRP tumor revealed that the somatic *FH:* c.1390+6T>A causes mis-splicing of 27.6% of the transcripts in tumor RNA, as indicated by the number of reads spanning consensus splice site (n=68) and the new cryptic splice site (n=26) (Figure 4B). None of the six carriers of predisposing variant in East Asian ancestry harbored a biallelic somatic mutation. Overall, the assessment of LOH and biallelic mutation supports the variants’ contribution to oncogenesis through the two-hit model.

**Figure 4.**
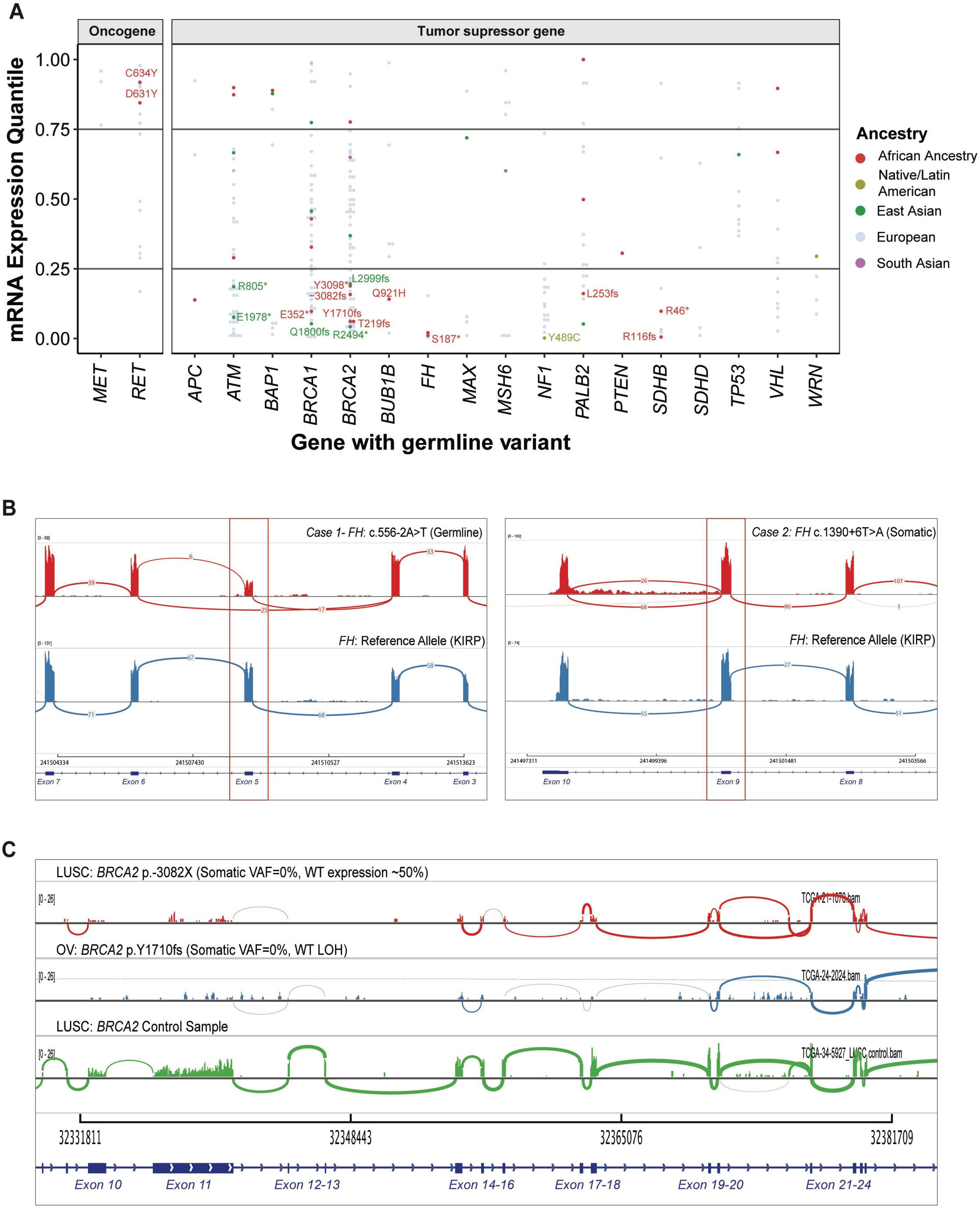
Expression changes associated with the predisposing variants. **A.** mRNA gene expression of the affected genes in the carriers of ancestry-specific variants as quantiles in their respective cancer cohort. Each dot denotes the gene expression level of a predisposing variant carrier colored by ancestry. Non-European variants corresponding to the bottom 25% expression in affected tumor suppressor genes and top 25% expression in affected oncogenes are further labeled. **B.** Plots of tumor RNAseq alignment highlighting exon (red box) with germline or somatic splice site variants in two cases with *FH* splice site variants as visualized using the integrated genome viewer (IGV). **C.** Tumor RNA expression for the *BRCA2* gene. The first two rows correspond to samples with a germline predisposing variant coupled with or without somatic LOH event, respectively. The third row corresponds to an unrelated sample without any *BRCA2* alteration. All three coverage plots are scaled together to show lower expression in the two samples harboring *BRCA2* alterations.

### Expression Changes in Predisposing Genes

To examine the transcriptional effects of the predisposing variants, we investigated the gene expression in tumor samples of the predisposing variant carriers (Figure 4A). We observed 154 overall and 27 non-European ancestry-specific predisposing variants cooccurring with an extreme expression (> 80% or < 20% in the same cancer cohort) of the respective gene, although the current sample sizes preclude us from discovering significantly-associated genes compared to non-carriers within each ancestry-cancer cohort (**Supp. Table 5a**).

All of the expression-associated variants were germline heterozygous variants at the DNA level. The degree of their variant allele fraction in the tumor RNAseq data (RNA VAF) thus indicates the degree of allelic-specific expression (ASE). The African carriers of *SDHB* truncating variants p.R116fs (the corresponding gene’s expression ranks at the bottom 0.5 percentile among all PCPG cases [0.5%], RNA VAF= 0.25 and p.R46* (9% in PCGP, RNA VAF= 0.80) showed low *SDHB* expression. The African carriers of *BRCA2* p.Y1710fs (6% in OV, RNA VAF =0) and p.3082fs (15% in LUSC, RNA VAF =0) also exhibited low *BRCA2* (Figure 4C). In the OV case, the germline *BRCA2* p.Y1710fs is coupled with a somatic LOH event, resulting in nearly complete loss of *BRCA2* expression.

Both of the African-ancestry carriers of FH predisposing variants, *FH* p.S187* (2% in KIRP, RNA VAF=0.13) and *FH*:c.556-2A>T (2% in KIRP, RNA VAF=0.50), showed low FH expression. In addition to the biallelic somatic *FH*:c.1390+6T>A mutation in the carrier of germline *FH* p.S187* described earlier, we also observed a mis-splicing event in a different case carrying germline *FH*:c.556-2A>T at the RNA level (Figure 4B).

For other ancestries, the tumor from one predisposing variant carrier of the Native/Latin American ancestry, *NF1* p.Y489C, showed low *NF1* mRNA expression (2% in BRCA, RNA VAF=0). Overall, RNA VAF of the majority of protein-truncating variants not accompanied by LOH varied between 0-0.25 (**Supp. Table 5a**), suggesting degradation of the mutant allele.

Many predisposing truncating variants of tumor suppressors are assumed to lead to loss of gene expression through mechanisms such as nonsense-mediated decay (NMD). Using the NMD Classifier[37], we revealed all frameshift variants found in the African and East Asian ancestries were located in the NMD-competent region (**Supp. Fig. 3**). These results support that a fraction of predisposing variants likely result in reduced gene products of tumor suppressors in ancestral groups.

Conversely for the rare tumors with germline variants in oncogenes, the two predisposing *RET* variants are coupled with elevated *RET* expression in their African ancestry carriers, including p.C631Y (84% in KIRC) and p.D634Y (91% in PCGP).

### Power Consideration for Predisposing Gene Discovery

Given the currently limited sample sizes in most of the minority cohorts, we sought to identify the required numbers of samples to discover novel cancer predisposing genes. We performed *post hoc* power analyses to detect a rare-variant association in an aggregation test using SKAT[24]. We assumed that a high proportion (80%) of variants are casual when focusing on prioritized predisposing variants in accordance with ACMG/AMP guidelines (**Supp. Table 6a**, see **Methods**)[15,16,22]. The detection of rare variants (AF<0.01) with moderate effect sizes (Odds Ratio [OR] >5) with at least 80% power requires sample sizes exceeding 1,000 samples (n=1014) per cancer type **(Supp. Figure 2A)**.

The sample size requirement suggests limited power for ancestry-specific analyses using TCGA, one of the largest cancer sequencing cohorts to date. For the largest ancestry subgroup in the study, European-ancestry BRCA cases (n=811), there is 67% power to detect genes with smaller effect sizes (OR < 3). For all other ancestries, their respective largest cohorts afford inadequate power to detect genes with large effect sizes (OR=9), including the African ancestry BRCA cohort (n=180, power=36%), the East Asian-ancestry LIHC cohort (n=162, power=24.5%), and the Native/Latin American-ancestry THCA cohort (n=11, power=<1%). As a reference, most known cancer predisposing genes, including *ATM, PTEN, STK11, CHEK2, BRIP1*, and *PALB2*, have an estimated OR < 10. *BRCA1/BRCA2* are exceptions with an OR > 10 for BRCA, but also show more moderate OR for other cancer types [38]. Despite limited power, this TCGA study includes three-fold more non-European cases (n=1,715) compared to the combined number of samples across 27 published non-TCGA sequencing studies that report race/ethnicity information from cancer cohorts (n=764 non-Europeans, 10 cancer types)[8]. Moreover, the majority of these studies focused on somatic alterations, and only a handful reported ancestry-specific germline predisposition (**Supp. Table 7**).

Standard power analyses have the caveat of assuming various unknown parameters that may be inaccurate. We thus performed a down-sampling analysis using two cancer types with at least five significantly associated germline genes in the European-ancestry: Pheochromocytoma and Paraganglioma (PCPG) and sarcoma (SARC)[3] (**Supp. Figure 2B, Supp. Table 6b**). We found that the sample size requirements differ for each gene and cancer cohort, likely due to varying penetrance. For example, six predisposing genes are discovered in both PCPG (n=146) and SARC (n=217) samples of the European ancestry, respectively, at their full cohort size. Upon down-sampling the cohort size in half, we found *VHL, SDHB, RET*, and *NF1* to be still associated in 73 PCPG cases, whereas only *TP53* remained significantly associated in 108 SARC cases. Even while assuming similar penetrance of the predisposing genes across ancestries, this analysis implicates that the discovery power is still far from saturation for most ancestry-specific cohorts (N < 100). The different predisposition landscapes across cancer types should also be accounted for in future study designs.

## DISCUSSION

We report one of the most extensive multi-ancestry investigations of rare cancer predisposing genes to date, encompassing 9,899 cancer cases across 33 cancer types. In the African ancestry, our results validated six known predisposing genes and nominated *BRCA2* as a potential predisposing gene for LUSC (Figure 1). In the East Asian ancestry, we unexpectedly found predisposing variants affecting *BRIP1* in STAD that warrants further investigation. Although the number of germline predisposing variants is small, they were associated with LOH (Figure 3), biallelic mutations (Figure 2), and gene expression effects in the tumor samples (Figure 4), supporting their potential contribution to cancer predisposition in carriers.

*Post hoc* power analyses highlight the need to expand each ancestral cancer cohort to at least 1,000 samples to confidently discover predisposing genes of intermediate or large effect sizes (OR > 5, **Supp. Figure 2**). It is necessary to use caution when interpreting the ancestry-specific predisposing gene associations identified herein or previous studies of smaller sample sizes, where a handful of carriers may give rise to the association in a limited cancer cohort. Further, the suggestive associations nominated by the TFT analyses will need to be established by analyses of larger cohorts adjusted for potential confounders. Two of the associations we identified in the African ancestry were also complemented by familial studies [27,30], providing further validation. To design future cancer genomics studies, one must note that the power considerations differ for discovering somatic driver genes and germline predisposing genes. Current detection powers have potentially reached saturation in detecting somatically mutated genes for sample sizes in multiple cancer types of TCGA[3], although racial disparities of the sequencing data could potentially limit the generalizability of findings [39–41]. We further highlighted the imbalanced dataset limits power for germline gene discovery in populations under-represented in research studies. In this TCGA cohort, we found multiple significant predisposing genes for the European ancestry and seven for the African ancestry, yet did not had a cancer cohort with sufficient testing samples for many other ancestries, including Native/Latin American and South Asian that each constitutes a considerable fraction of the US population.

We observed selected predisposing genes shared across ancestries (ex. *BRCA2* in BRCA/OV and *SDHB* in PCPG for both the African and European ancestries). Predisposing variants, on the other hand, are highly ancestry-specific (Figure 2). Many of the predisposing variants found in the African or East Asian ancestry were not identified in the much larger European-ancestry population of TCGA (n=8,184) or even the gnomAD non-cancer cohort (n=134,187) or submitted to ClinVar by clinical laboratories assessing patients for cancer predisposition. Rare variant classification and interpretation remain a challenge given the low frequency of observation precluding statistical associations. The identification of ancestry-specific predisposing variants further highlights this challenge in minority groups, where current germline sequencing often results in higher rates of variants of unknown significance (VUSs)[10].

Personalized medicine provides tailored disease diagnosis and treatment plans based on an individual’s unique genetic profile. The knowledge of different cancer predisposing genes and prevalence across ancestries suggests that we need to provide ancestry-specific interpretations of genetic data. In particular, many of the current guidelines for when genetic testing is recommended rely on the underlying likelihood of identifying a germline variant. Thus, accurate estimates of germline prevalence may alter recommendations for different patient populations. At the current sample sizes for minority cohorts, our study is still limited in power to discover and establish ancestry-specificity of predisposing genes (**Supp. Figure 2**). However, we we were able to discover many ancestry-specific variants not currently submitted to ClinVar. Further, much of the diverse populations within the US, not to mention world-wide, still lack representation in existing sequencing cohorts. Ongoing sequencing projects will begin to address this disparity within US populations and multiple countries in East Asia and Europe [42]. Yet, many populations, such as the diverse African ancestry [43], remain underserved although projects like H3Africa are designed to address this problem. Additional efforts will be required to deliver the promise of genome-based precision medicine for all.

To aid interpretation of low-frequency ancestry-specific variants, evidence of a somatic second hit event (i.e., loss of heterozygosity [LOH] or a biallelic mutation) in tumor samples can support functionality. Our analysis of the two-hit model identified the second somatic events in two thirds (10/15) of the African-ancestry specific predisposing variants and in one out of six of the East Asian ancestry-specific predisposing variants (**Supp. Table 4b**). Additionally, some carriers of ancestry-specific predisposing variants showed simultaneous extreme expression of the affected genes (Figure 3). Such evidence derived from analysis of the somatic genome or transcriptome can be further utilized to characterizing rare germline variants[44], especially when DNA-level analysis still suffers from limited sample sizes.

Our observation of somatic second hit (Figure 2-3) and transcriptional effects (Figure 4) coupled with germline variants also adds on to the current literature on germline-somatic interactions in cancer [45]. While the majority of cancer genomic studies focus exclusively on the germline or somatic genome, pathogenic germline variants are associated with different somatic mutational signatures, allele-specific imbalance, or somatic drivers [11,26,46–48]. The availability of germline DNA analysis and tumor genomic and transcriptomic analyses from the same individual provides critical data to the analyses describe here that is not possible in many studies that only analyze germline DNA samples alone. Collectively, these findings are providing the roadmaps of how germline variants may trigger and collaborate with specific somatic mutations, eventually leading to cancer development. In this process, genomes across different ancestral populations provide different contexts for developing somatic mutations and genomic instability, even when the individual carries the same germline predisposition variant. We showcased examples of predisposition-associated LOH and gene expression changes in diverse individuals. As sample sizes of sequencing cohorts expand, analyzing germline-somatic interactions across ancestry will be pivotal to reveal potential ancestry-specific effects.

In summary, we identify ancestry-specific predisposing genes and variants contributing to multiple cancer types. The results provide insights into rare genetic predisposition and their somatic impacts in cases of African and East Asian ancestries. Continuous studies using larger ancestry cohorts will be required to enable adequately-powered discovery of predisposing genes, which will, in turn, improve genetic screening and diagnostic strategies for diverse populations.

## DECLARATIONS

### AUTHORS’ CONTRIBUTIONS

KH and NO conceived and designed the research and analyses. NO, KH, RB, SEP, and LD acquired the data. The PanCanAtlas AIM working group, AC, JM, and KH conducted the genetic ancestry assignment. NO and KH conducted the analyses. NO and KH interpreted the results and drafted the manuscript. KH supervised the study. All authors read, edited, and approved the manuscript.

## ACKNOWLEDGEMENTS

The authors wish to acknowledge The Cancer Genome Atlas and its participating patients and family that generously contributed the data. The authors would also like to acknowledge members of TCGA PanCanAtlas Research network, particularly active members of the Germline Analysis Working Group and the Ancestry Information Markers Analysis Working Group, for helpful discussions. ZHG acknowledges funds from the LUNGevity Foundation.

### ^@^TCGA Analysis Network

Jian Carrot-Zhang^1,2^, Nyasha Chambwe^3^, Jeffrey S. Damrauer^4^, Theo A. Knijnenburg^3^, A. Gordon Robertson^5^, Christina Yau^6,7^, Wanding Zhou^8^, Ashton C. Berger^1,2^, Kuanlin Huang^9^, R. Jay Mashl^10^, Justin Newberg^11^, Alessandro Romanel^12^, Rosalyn W. Sayaman^13,14^, Francesca Demichelis^12^, Ina Felau^15^, Garret Frampton^11^, Seunghun Han^1,2^, Katherine A. Hoadley^4^, Anab Kemal^15^, Peter W. Laird^8^, Alexander J. Lazar^16^, Xiuning Le^17^, Ninad Oak^18, 19^, Hui Shen^8^, Christopher K. Wong^20^, Jean C. Zenklusen^15^, Elad Ziv^13,14^, Francois Aguet^1^, Li Ding^6^, John A. Demchok^15^, Michael K.A. Mensah^15^, Roy Tarnuzzer^15^, Zhining Wang^15^, Liming Yang^15^, Jessica Alfoldi^1^, Konrad J. Karczewski^1^, Daniel G. MacArthur^1^, Garret M. Frampton^11^, Christopher Benz^6^, Joshua M. Stuart^20^, Andrew D. Cherniack^1,2^, Rameen Beroukhim^1,2,21^

1. The Eli and Edythe L. Broad Institute of Massachusetts Institute of Technology and Harvard University, Cambridge, MA 02142, USA

2. Department of Medical Oncology, Dana-Farber Cancer Institute, Boston, MA 02215, USA

3. Institute for Systems Biology, Seattle, WA 98109, USA

4. Lineberger Comprehensive Cancer Center, University of North Carolina at Chapel Hill, Chapel Hill, NC 27599, USA

5. British Columbia Cancer Agency, Genome Sciences Centre, Vancouver, Canada V5Z 4S6

6. Buck Institute for Research on Aging, Novato, CA 94945, USA

7. Department of Surgery, University of California, San Francisco, San Francisco, CA 94115, USA

8. Van Andel Research Institute, Grand Rapids, MI 49503, USA

9. Department of Genetics and Genomics, Icahn School of Medicine at Mount Sinai, New York, NY 2129, USA

10. Department of Medicine, Washington University in St. Louis, St. Louis, MO 63110, USA

11. Cancer Genomics Research, Foundation Medicine, Inc., Cambridge, MA 02141, USA

12. Department of Cellular, Computational and Integrative Biology (CIBIO), University of Trento, Via Sommarive 9 Povo (TN) 38123 Italy

13. Department of Laboratory Medicine, Helen Diller Family Comprehensive Cancer Center, University of California San Francisco, San Francisco, CA 94143, USA

14. Department of Population Sciences, Beckman Research Institute, City of Hope, Duarte, CA 9210

15. National Cancer Institute, Bethesda, MD 20892, USA

16. Departments of Pathology, Genomic Medicine, and Translational Molecular Pathology, The University of Texas M.D. Anderson Cancer Center, Houston, TX 77030, USA

17. Department of Thoracic and Head and Neck Medical Oncology, The University of Texas M.D. Anderson Cancer Center, Houston, TX 77030, USA

18. Department of Oncology, St. Jude Children’s Research Hospital, Memphis, TN 38105, USA

19. Department of Molecular and Human Genetics, Baylor College of Medicine, Houston, TX 77030, USA

20. Department of Biomolecular Engineering, Center for Biomolecular Sciences and Engineering, University of California, Santa Cruz, Santa Cruz, CA 95064, USA

21. Department of Medicine, Brigham and Women's Hospital and Harvard Medical School, Boston, MA, USA

## ADDITIONAL FILES

### Supplementary Figures

**Supp. Figure 1.** Principal component analyses (PCA) of germline TCGA samples to infer genetic ancestry as performed by PanCanAtlas Ancestry Informative Markers (AIM) working group

**Supp. Figure 2.** Power analysis for ancestry-specific sample sizes to discover predisposing genes.

**A.** The number of samples required to detect rare variant associations with varying effect sizes (OR = 2,3,5,7,9) at corresponding statistical power. The total number of samples assumes an equal number of cases and controls (e.g. For 811 TCGA-BRCA- EUR samples, n=1,622). Cancer types with the largest cohort sizes for each of the studied ancestries in TCGA are shown by a dotted line.

**B.** Down-sampling analysis to identify counts of significantly associated predisposing genes at different sample sizes by incrementally increasing the sample size from zero to the current cohort sizes.

**Supp. Figure 3.** Nonsense-mediated decay prediction for predisposing frameshift variants in African and East Asian ancestries

### Supplementary Tables

**Supp. Table 1.** The demographic information of TCGA PanCanAtlas cohort with separate admixture populations

**Supp. Table 2a.** Ancestry-specific cancer-gene associations discovered from multivariate regression analyses.

**Supp. Table 2b.** Ancestry-specific cancer-gene associations discovered from rare variant burden testing (Total Frequency Test- TFT).

**Supp. Table 3.** Frequency of predisposing variants in TCGA PanCanAtlas and gnomAD-non-cancer subset across all ancestries.

**Supp. Table 4a.** Ancestry-Specific Predisposing Variants as identified from Supp. Table.2

**Supp. Table 4b.** Summary of somatic second hit mutations in carriers of germline predisposing variants.

**Supp. Table 5a.** Statistical analysis of gene expression in tumor samples of the variant carriers vs. non-carriers within each ancestry-cancer combination

**Supp. Table 5b.** Tumor RNAseq variant allele fractions and the somatic second hit events in germline predisposing variants with extreme expression within that cancer type.

**Supp. Table 6a.** *Post hoc* power analyses to detect rare-variant associations in an aggregation test using SKAT.

**Supp. Table 6a.** Down-sampling analysis for PCGP and SARC (cancers with at least 5 significantly associated germline genes in the European ancestry).

**Supp. Table 7.** Prior studies that report ancestry-specific germline predisposition.

