## Supplementary Figures for "Ancestry-Specific Predisposing Germline Variants in Cancer"

**Supp. Figure 1** Principal component analyses (PCA) of germline TCGA samples to infer genetic ancestry as performed by PanCanAtlas Ancestry Informative Markers (AIM) working group

UCSF PCA

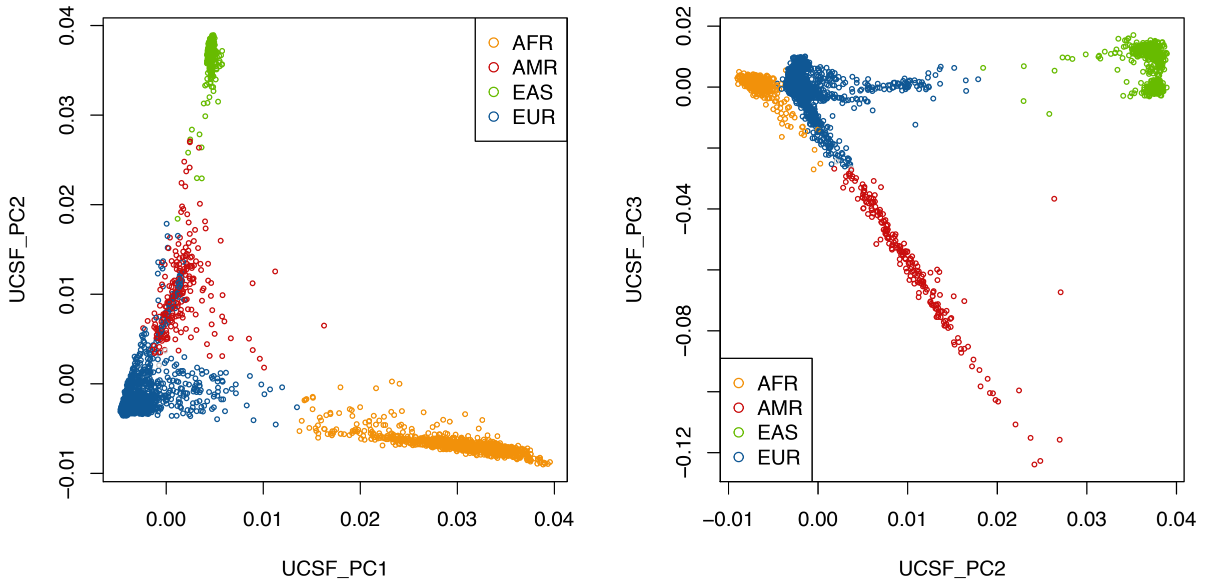


**WashU PCA**

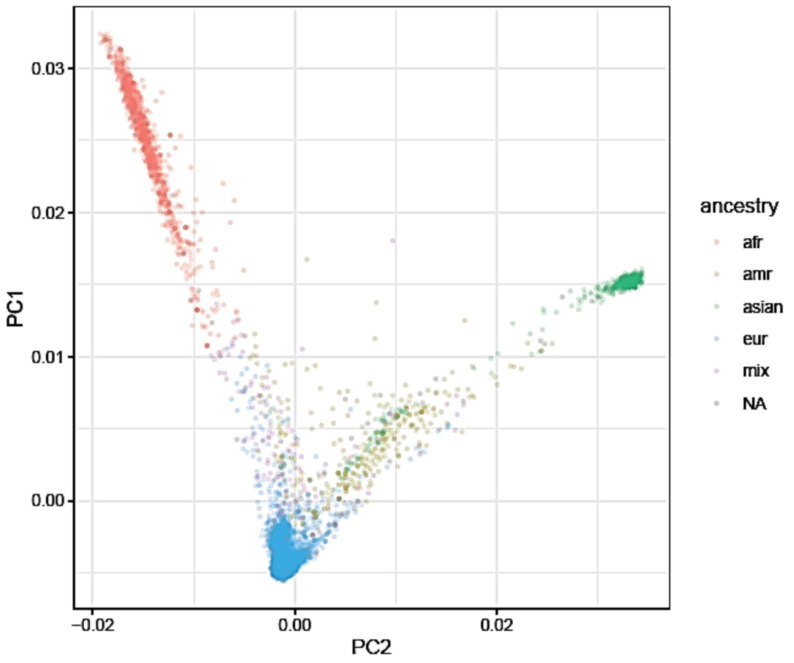

**Supp. Figure 2** Power analysis for ancestry-specific sample sizes to discover predisposition genes

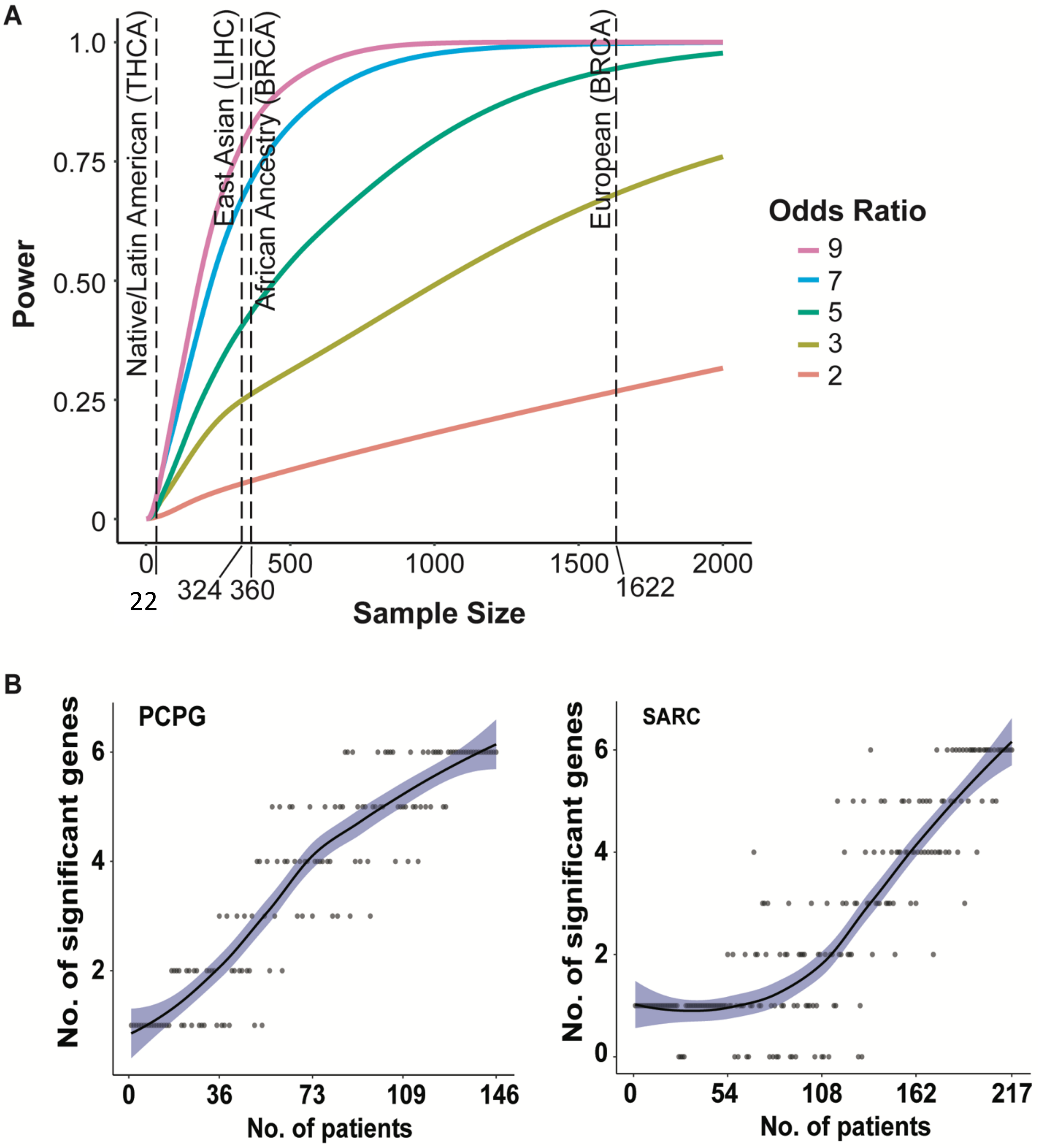

**Supp. Figure 3** Nonsense-mediated decay prediction for predisposing frameshift variants in African and Asian ancestries

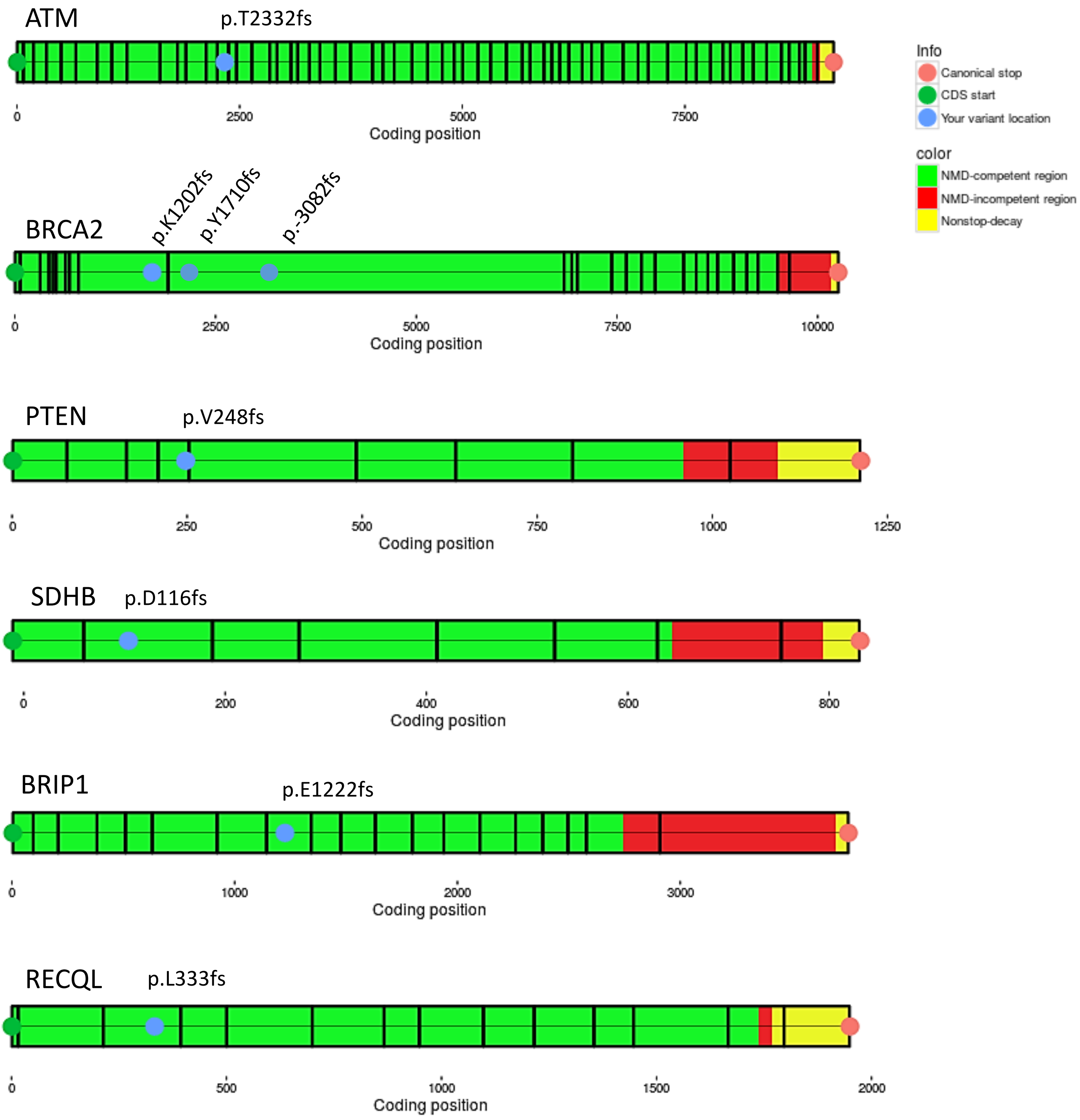
